# Particle-based simulation reveals macromolecular crowding effects on the Michaelis-Menten mechanism

**DOI:** 10.1101/429316

**Authors:** Daniel R. Weilandt, Vassily Hatzimanikatis

## Abstract

Many computational models for analyzing and predicting cell physiology rely on *in vitro* data, collected in dilute and cleanly controlled buffer solutions. However, this can mislead models because about 40% of the intracellular volume is occupied by a dense mixture of proteins, lipids, polysaccharides, RNA, and DNA. These intracellular macromolecules interact with enzymes and their reactants and affect the kinetics of biochemical reactions, making *in vivo* reactions considerably more complex than the *in vitro* data indicates. In this work, we present a new type of kinetics that captures and quantifies the effect of volume exclusion and any other spatial phenomena on the kinetics of elementary reactions. We further developed a framework that allows for the efficient parameterization of this type of kinetics using particle simulations. Our formulation, entitled GEneralized Elementary Kinetics (GEEK), can be used to analyze and predict the effect of intracellular crowding on enzymatic reactions and was herein applied to investigate the influence of crowding on phosphoglycerate mutase in *Escherichia coli*, which exhibits prototypical reversible Michaelis-Menten kinetics. Current research indicates that many enzymes are reaction limited and not diffusion limited, and our results suggest that the influence of fractal diffusion is minimal for these reaction-limited enzymes. Instead, increased association rates and decreased dissociation rates lead to a strong decrease in the effective maximal velocities *V_max_* and the effective Michaelis-Menten constants *K_M_* under physiologically relevant volume occupancies. Finally, the effects of crowding in the context of a linear pathway were explored, with the finding that crowding can have a redistributing effect, relative to ideal conditions, on the effective flux responses in the case of two-fold enzyme overexpression. We suggest that the presented framework in combination with detailed kinetics models will improve our understanding of enzyme reaction networks under non-ideal conditions.

## Introduction

The intracellular environment is a crowded place, with about 20–40% of the interior volume of living cells occupied by a variety of macromolecules including proteins, RNA, DNA, and lipids. (1, 2). The composition of this mixture depends considerably on the organism and its environment, but even within the cell, the local density and size distribution of the macromolecules varies between and within compartments (3–5). Because the presence of these macromolecules can impact diffusion rates, protein conformation, folding and aggregation, catalytic rates, and enzyme-substrate affinities (6–8), an alteration in the elementary properties governing the spatiotemporal dynamics of cells can affect all cellular functions, such as expression, translation signaling, and metabolism. Because many of these cellular functions depend on specific reactions catalyzed by cellular enzymes, it is necessary to study the effect of macromolecular crowding on the function of enzyme-catalyzed reaction systems, and to do this, it is necessary to characterize the kinetics of these systems under the altered conditions.

Computational models are used to analyze and predict cell physiology, though computational studies are limited in their frequent reliance on *in vitro* characteristics to directly parameterize their models (9, 10), reduce uncertainty (11, 12), or to evaluate predicted parameters (13). This causes that the actual enzyme *in vivo* characteristics are not captured such that the model predictions from these studies might deviate significantly from the ones measured *in vitro*. This is especially true considering that *in vitro* characterizations are usually performed in dilute, homogenous conditions whereas reactions in the cytoplasm occur in an inhomogeneous and densely packed environment (14).

The importance of environmental impact on enzyme kinetics is therefore an important topic of study, especially in terms of crowding in the densely packed intracellular space. In early studies of crowded enzyme catalysis, it was believed that the main effect of diffusion-limited Michaelis-Menten kinetics was caused by altered, anomalous diffusion accompanied by increased effective concentrations. These studies were limited, however, in that volume exclusion effects caused by the reactive partners themselves were often neglected, which results that the change in activity due to interaction with macromolecules is not captured 15–18). A more recent work on the subject was presented by Mourao *et al*. in which fractal behavior, indicating that the diffusion and the apparent order of the elementary reactions is altered, was studied using a lattice-based model for an irreversible Michaelis-Menten mechanism. They showed that fractal kinetics only occur under very restrictive conditions, suggesting that it might be less common than previously assumed (19).

Further recent work has shown that the effective rate constant for bimolecular reactions changes under crowded conditions (20, 21). Berezhkovskii and Szabo demonstrated that it is possible to express the effective rate for bimolecular reactions as a function of a *crowding-induced interaction potential* between two reaction partners, which results from an interaction with the surrounding particles when two reactants are in contact. Repulsive interactions between the reactants and particles would, therefore, result in an attractive effective potential between the reactants and vice versa. Relatedly, it has been shown that for rate-limited reactions in the presence of high reactant concentrations, the influence of diffusion minimal, indicating that the effective *crowding-induced interaction potential* might be more dominant for most enzymatic reactions (22).

Because of its importance in modeling *in vivo* systems, the effects of crowding on biochemical reactions have been extensively studied by various computational and experimental methods, as seen in several reviews (1, 2, 23). Most of the effort in these studies has been directed towards investigating the impact of diffusion in fractal media on the reaction kinetics (22), with little focus on characterizing the effect of crowding on the mean effective enzyme kinetics. However, since it has now been shown that most enzymes are not diffusion limited but instead are reaction limited (24), the reevaluation of crowding in these reactions is important. In this work, therefore, we aimed to design computational methods for studying spatial effects of any kind, applying our work specifically to the effects of crowding on reaction-limited instead of diffusion-limited enzymes, with a goal of bridging the discrepancy between the *in vitro* measurement of kinetic parameters and the actual *in vivo* conditions.

In contrast to the previous studies on the Michaelis-Menten kinetics, that use a diffusion limited irreverisble reaction scheme, we studied the effect of crowding on enzyme kinetics by employing a fully reversible reaction scheme and present herein an example with a representative catalytic activity and affinities that result in a reaction-limited enzyme. Additionally, our molecular particle model accounts for volume exclusion and the diffusion of all species, including reactants and crowding agents, and this was used to study the effect of different size distributions of crowding agents on reaction kinetics.

Previous studies into crowding conditions are limited by their computational cost and lack of global insight into the sensitivity of the reaction kinetics. They often use spatial simulation techniques to simulate multiple realizations of reaction trajectories to determine the influence on the effective kinetics under very specific conditions, meaning that these studies only gain insight into the local sensitivity of the kinetics with respect to the crowding conditions. Furthermore, it is computationally very expensive to resolve the reaction trajectories from particle simulations for reaction-limited reactions because the time scale to resolve the diffusion of the particle is up to seven orders of magnitude faster than the reaction time scale. This requires billions of time steps to be solved for tens of thousands of particles, resulting in a month of simulation time for a single trajectory (25).

To resolve these challenges, we present a new formulation entitled GEneralized Elementary Kinetics (GEEK) to characterize the kinetic mechanisms that are influenced by various spatial effects, including volume exclusion, confinement (1D/2D diffusion), strong and weak interaction forces, localization, or any combination of similar phenomena. In the presented work, we use a coarse-grained particle model based on hard-sphere Brownian reaction dynamics to parametrize this formulation, which can be used in a straightforward way to build ordinary differential equation (ODE) models that use power-law approximations to capture the characteristics of spatial effects and to directly quantify the impact of fractal diffusion.

We applied our method to the investigation of macromolecular crowding on the function of phosphoglycerate mutase (PGM) in *Escherichia coli*. Our presented example clearly demonstrates that accounting solely for an increased local concentration and anomalous diffusion is not sufficient to properly describe crowding effects. We show that a mechanism-dependent effect emerges upon crowding that is facilitated by an increase in both product and substrate association activity and a decrease in the dissociation activity. For reversible Michaelis-Menten kinetics, these effects result in an increase in the binding affinity for the product and substrate as well as a decrease in the maximal reaction rate. Finally, we investigated the effects of crowding on a linear pathway, where we show that crowding can significantly redistribute the relative flux responses with respect to enzyme overexpression, indicating that the impact of altered kinetics is also propagated on a network level.

## Methods

### Reversible Michaelis-Menten kinetics

In this study, we primarily investigated a reversible Michaelis-Menten reaction mechanism, where a substrate *S* binds to an enzyme *E* to form a complex *ES* via a reversible reaction, which can reversibly transform the substrate and reversibly dissociate the product *P*. The overall reaction scheme is given by:

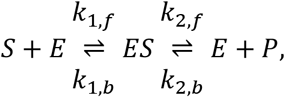

where *k*_1,*f*_, *k*_1,_*_b_*, *k*_2,_*_f_*, and *k*_2,*b*_ denote the rate constants of the elementary reactions. The typical form of the reaction rate *v* as a function of substrate and product concentration, see Eq. (2), is derived from the assumption that all enzymes are conserved such that [*ES*] + [*E*] = [*E_T_*], where [*E_T_*] denoted the total enzyme concentration, and the enzyme-substrate complex concentration [*ES*] is in a quasisteady state, i.e. *d*[*ES*]/*dt* ≈ 0 (26). 
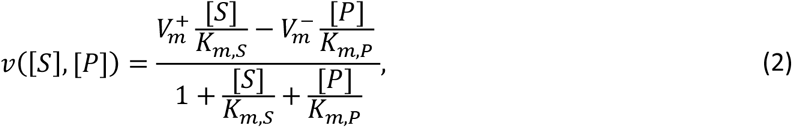
where the parameters 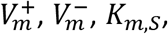 and *K_m,P_* are related to the elementary rate constants *k*_1/2,*f/b*_ given a [*E_T_*]. 
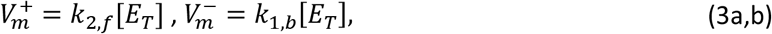

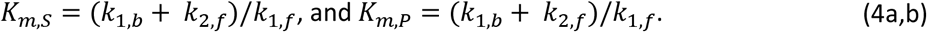

The equilibrium constant of the system is then: 
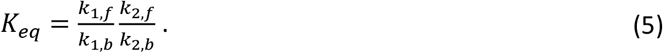

### Generalized elementary kinetics (GEEK)

By introducing inert molecules, we observe an alteration of the effective rate constants due to a change in the diffusion and the collision dynamics. In the most general case, this can, compared mass-action kinetics, result in a change of the effective order and effective rate constant. Berezhkovskii and Szabo showed that the effective (Collins-Kimball) reaction rate constant *k_CK_* for a diffusion-influenced, bimolecular reaction under crowded conditions can be expressed in terms of an altered diffusion constant *D_1_* and an external *crowding induced interaction potential* Δ*U* between the two reacting species. This potential is an implicit representation of the interaction of the individual reactant species with the molecules in their environment and whether these interactions keep the reactants in contact or if they are tearing them apart. The expression for the Collins-Kimball rate constant was found to follow that described by Berezhkovskii and Szabo (20): 
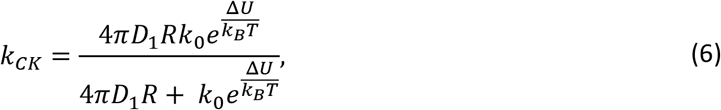
where *k*_0_ is the reaction rate upon collision. For the reaction-limited case *k*_0_ ≫ 4n*D*_1_*R*, this expression simplifies to an exponential relation, *k_CK_* ≈ *k*_0_ exp(Δ*U/k_B_T)*. In general, the induced interaction potential *ΔU* and the diffusion constant *D*_1_ are a function of the global state of the system, which includes concentrations and intermolecular interactions.

To approximate this deviation of the effective elementary rate constants, indicated by *k_j,eff,_*, where j *∊* [(1,*f*), (1,*b*), (2,*f*), (2,*b*)], from the free rate constants in ideal conditions, *k_j,0_* in a general form, the logarithmic deviation *ζ_j_* was introduced: 
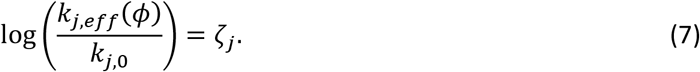

To quantify this deviation as a function of the species’ concentrations, a linear function of the scaled logarithmic concentrations log([*X_i_*]/[*X_i_*]_0_) was introduced and tested for *ζ_j_*:

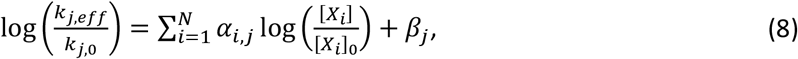

where *α_i,j_* are the coefficients quantifying the effect of one of the *N* reactants *X_i_* on reaction j, and *β_j_* is an offset attributed to the effect of different occupied volume fractions.

The effective reaction rate is thus given by:

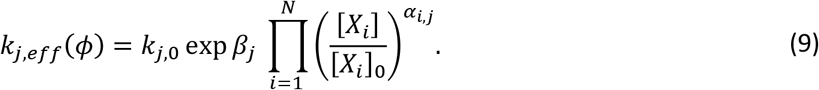

From this expression, a generalized mass-action rate law is defined for the elementary reactions:

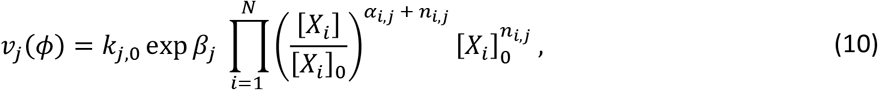

where *n_i,j_* denotes the stoichiometric coefficient of the substrate *X_i_* in reaction *j*. The generalized elementary mass-action rate law (10) can be directly used to create a system of ordinary differential equations that can approximate the time evolution of the system under non-ideal conditions.

### Generalized elementary Michaelis-Menten kinetics

Given the generalized elementary rate laws, the quasi-steady-state approximation for the Michaelis-Menten reaction rate with generalized elementary kinetics can be defined. Therefore, it can be assumed that the enzyme is conserved, [*ES*] + [*E*] = [*E_T_*], and the enzyme-complex is in a quasisteady state, i.e.

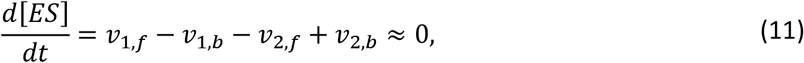

where each flux *V_j_* given by a generalized rate law as given in Eq. (10). The reaction rate of the enzymatic reaction at steady state is then given by the rate of product formation at steady state:

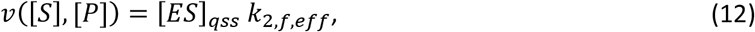

where [*ES*]_*qss*_ is enzyme complex concentration at the quasi-steady state. For the case of 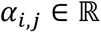 and *α_i,j_* ≠ 0, it is not possible to obtain an explicit expression for the reaction rate *v*([*S*], [*P*]). Eq. (11) can thus be solved numerically for the quasi-steady-state concentration of the enzyme complex [*ES*]_*qss*_ for given amount of total enzyme [*E_T_*], substrate [*S*], and product [*P*], and the reaction rate can then be calculated according to Eq. (11). The average apparent Michaelis-Menten parameters are then extracted using a linear approximation of *v*([*X*]) with *v*([*X*])/[*X*] for either [*S*] = 0 or [*P*] = 0, i.e. the Eadie-Hofstee form of Michaelis-Menten kinetics (27, 28). The slope of these linear regressions yields the respective *K_m_*, and the y-axis intercept yields the respective *V_max_*:

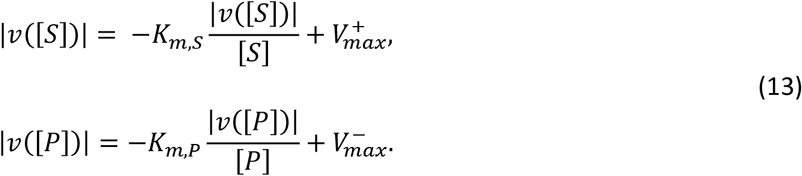

To express the thermodynamic driving forces, the elementary rate model was considered as *M* reversible reactions *ρ ∊* [1,2], with the forward flux *v_ρ,f_* and the backward flux *v_ρ,b_*. Using the principle of detailed balance, the free energy of the reaction can be expressed as a function of the displacement from equilibrium Γ = *v_b_*/*V_f_* (29):

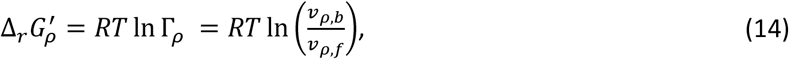

where *R* is the general gas constant and *T* is the absolute temperature. With the fluxes expressed in terms of the generalized elementary rate law (10), the free energy reads:

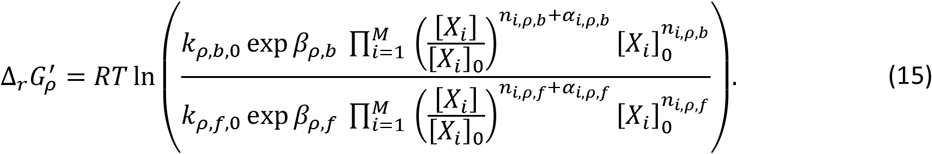

In general, the overall free energy consists of the ideal and non-ideal contributions. The ideal contribution consists of the standard free energy of the reaction and the concentration contributions, and the non-ideal contribution contains terms emerging from molecular interactions, such as by steric repulsion, van der Waals forces, and electrostatic interactions.

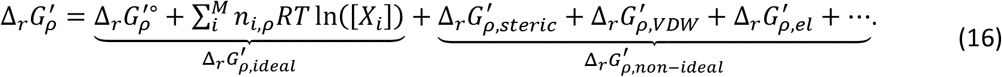

In this work, only the non-ideal contributions due to steric repulsion were modeled by means of a hard-sphere potential, though in the most general case, the formulation presented in this work allows for the inclusion of any kind non-ideal contributions:

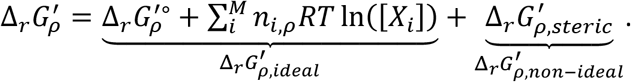

By comparing the free energy of the generalized elementary rate model to the free energy of the dilute mass-action equivalent, the ideal contribution can be identified as:

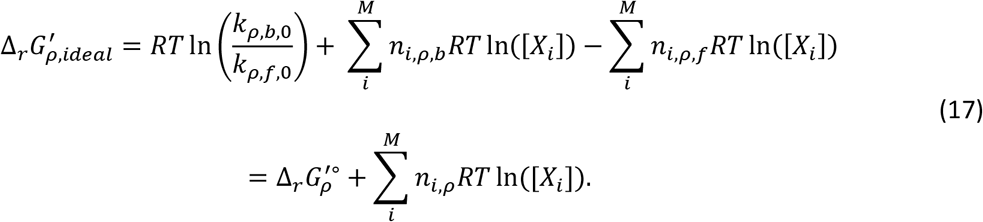

The remaining contributions can be identified as the non-ideal contribution:

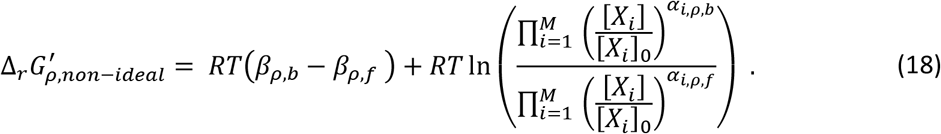

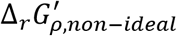 can be further partitioned into a reactant-independent and a reactant-dependent contribution:

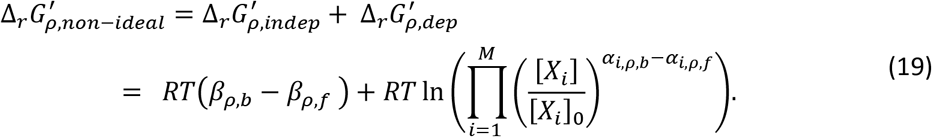

The free energy of the generalized elementary Michaelis-Menten kinetics is given by the sum of all reversible reaction free energy contributions 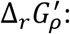

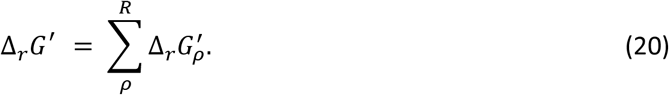

With [*X_i_*] = [[*S*], [*E*], [*ES*], [*P*]], the free energy of the reaction can be simplified to the well-known ideal contribution containing only the chemically modified species [*S*] and [*P*] as well as a non-ideal contribution, wherein the non-ideal contribution is a phenomenological description of free energy change based on the generalized elementary kinetics.

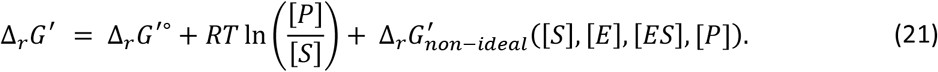

The formulation of the reversible Michaelis-Menten rate law in terms of generalized elementary kinetics allows for the phenomenological capture of non-linear effects on the collision level.

### Hard-sphere Brownian reaction dynamics

To incorporate the spatial effects into the enzymatic reaction system, hard-sphere Brownian reaction dynamics (HSBRD) were used. The method used in this work is a straightforward combination of hard-sphere Brownian dynamics (30) and Brownian reaction dynamics (31, 32) that was accomplished by adding the hard-sphere collision dynamics as described to Brownian reaction dynamics. The method describes the movement of independent particles as a random walk of point particles diffusing in a viscous medium. Thereby, HSBRD neglects the hydrodynamic interactions between the particles. The equations of motion are given in terms of the overdamped Langevin equation. Using the Einstein-Smoluchwoski relation, its velocity is given by Wang and Uhlenbeck (33):

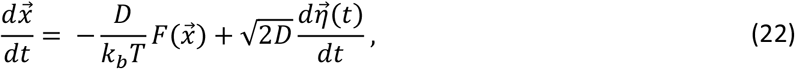

where 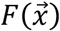 is a force acting on the particle, *k_b_* is the Boltzmann constant, *T* is the absolute temperature of the surrounding fluid, and 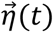 is the result of a three-dimensional Wiener process. An explicit Euler formulation was used to update the positions at every time step, *Δt*, as follows:

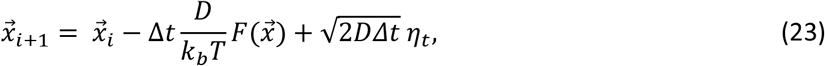

where *η_t_* is a random vector drawn from a normal distribution.

When two reactants collided, i.e. their radii overlapped after the positions were updated, a uniform distributed random number *r* was compared to the reaction with a probability *p* to determine if the reaction occurred within this time step Δ*t*. The probability *p* was determined by a microscopic reaction rate *k_micr_* (34):

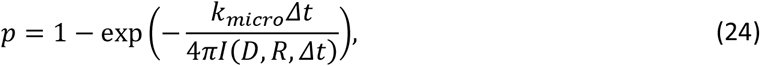

where *I*(*D,R,Δt*) is a normalization factor for the effective collision volume in Brownian reaction dynamics simulations, as derived by Morelli and ten Wolde (34). The observed steady-state reaction rate *k_AB_* for a bimolecular reaction in homogenous, dilute conditions is then given by (35):

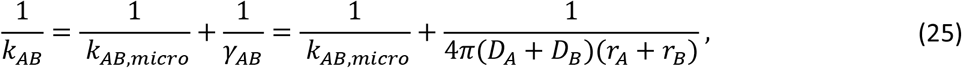

where *k_AB,micro_* is the reaction rate upon collision, and *γ_AB_* is the diffusion-limited reaction rate constant.

In the case of a particle collision without a subsequent reaction, an elastic hard-sphere collision was assumed to take place. The new particle position was computed from the momentum conservation using the average velocity 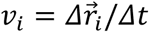 of the move that led to the particle overlap (30).

First-order reactions are modeled similarly to bimolecular reactions by comparing a uniformly distributed random variable to the probability that the reaction took place in the time interval Δ*t*, with the reaction probability of *p* = 1 — exp(−*k_micro_Δt*). The reaction products are placed in contact around the original position of the educt using a random orientation. If the products were to collide with any other particles, the move would be rejected and the educt would remain at its original position. Otherwise, the educt would be removed and the products would be placed instead.

Furthermore, constant particle boundary conditions were applied at every timestep through the random insertion or removal of particles of a given species to match the specified particle count of the species. The HSBRD particle simulation was implemented in C++ using the OPENFPM framework (http://openfpm.mpi-cbg.de/).

### Measuring effective rate constants

Since it is necessary to resolve the particle movement on the nanosecond timescale as opposed to the timescales of the reaction dynamics, which are found to be on the order of seconds to hours, a separated timescale approach was proposed to efficiently bridge these differences. The effective elementary reaction rates at constant concentrations and crowding conditions in particle simulation were therefore measured. To measure the effective rate constants from a particle simulation, two separate schemes for monomolecular and bimolecular rate constants were proposed.

For monomolecular rate constants, the effective reaction rates were extracted by probing the space around the enzyme-substrate complexes. Therefore, every valid educt *k* for a reaction *j* was selected, and multiple dissociation reactions were attempted in a random direction. If the dissociation were to be successful, meaning that the dissociated particles did not collide, *ω_jlj_* = 1. If the dissociation were to yield a collision, *ω_jkl_* = 0. Averaging over the results of all dissociation attempts *ω_ij_* of the probed particle *i*, a local success probability of 〈*ω_kj_*〉 was obtained. To describe the equivalent homogenous system, the success probability 〈*ω_j_*〉 was computed as the mean of the local average success rate over all probed particles. In the limit of continuous concentrations, the effective rate constant, *k_eff,j_*, is given by the rate constant *k_0,j_* scaled by the success probability 〈*ω_j_*〉:

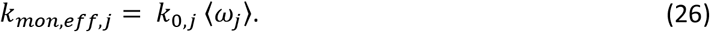

The effective bimolecular rate constant can be extracted from the effective collision frequency *Z_A,B_* between two species, A and B. This collision frequency is estimated as the number of collisions between A and B in an integrated time interval *C_A,B_*(*t, t* + *Δt*) per time step *Δt:*

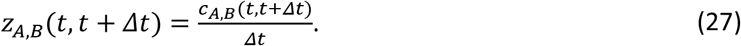

Given the probability of a reactive collision (Eq. (24)), the effective bimolecular rate constant can be measured as the mean collision frequency per number of possible interactions pairs, i.e. *N_A_N_B_*, scaled by the fraction of successful collisions within a measurement time interval *Δt:*

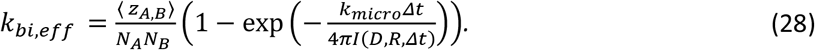

### Modeling Framework

In this paper, we propose a new simulation framework using the above-described concept of generalized elementary kinetics. In our simulation framework, an equivalent particle model is first created from an elementary step mechanism (Fig. 2, part 1). To create an equivalent particle model, only the elementary reactions of the enzyme mechanism are required. If only phenomenological constants, e.g. parameters for the quasi-steady-state approximation, are given for the enzymatic reaction, it is necessary to map these to the elementary reaction rate constants. Furthermore, all species involved in the elementary reaction properties need to be assigned to describe their molecular movement, i.e., a diffusion coefficient, a collision radius, and a mass. Given the molecular data, the rate constants of the particle model are matched with the rate constants of the elementary step model with the assumption that the measured, or calculated, rate constants were measured in homogenous, dilute conditions. In the case of monomolecular reactions, the observed rate constants are then equivalent to the microscopic transition rates. For bimolecular reactions, the diffusion-limited rate constant *γ_A,B_* is first computed based on the diffusion coefficients and collision radii and then matched to the effective reaction rate of the dilute, homogenous particle system with the rate constant in the elementary step model by adapting the corresponding microscopic rate constant *k_A,B,mic_* using Eq. (25). A volume that is large enough to capture the local bulk properties of a locally well-mixed enzyme substrate system is then chosen such that the number of particles of each species in the system is large enough to discretize the concentration space of interest.

**Figure 1:**
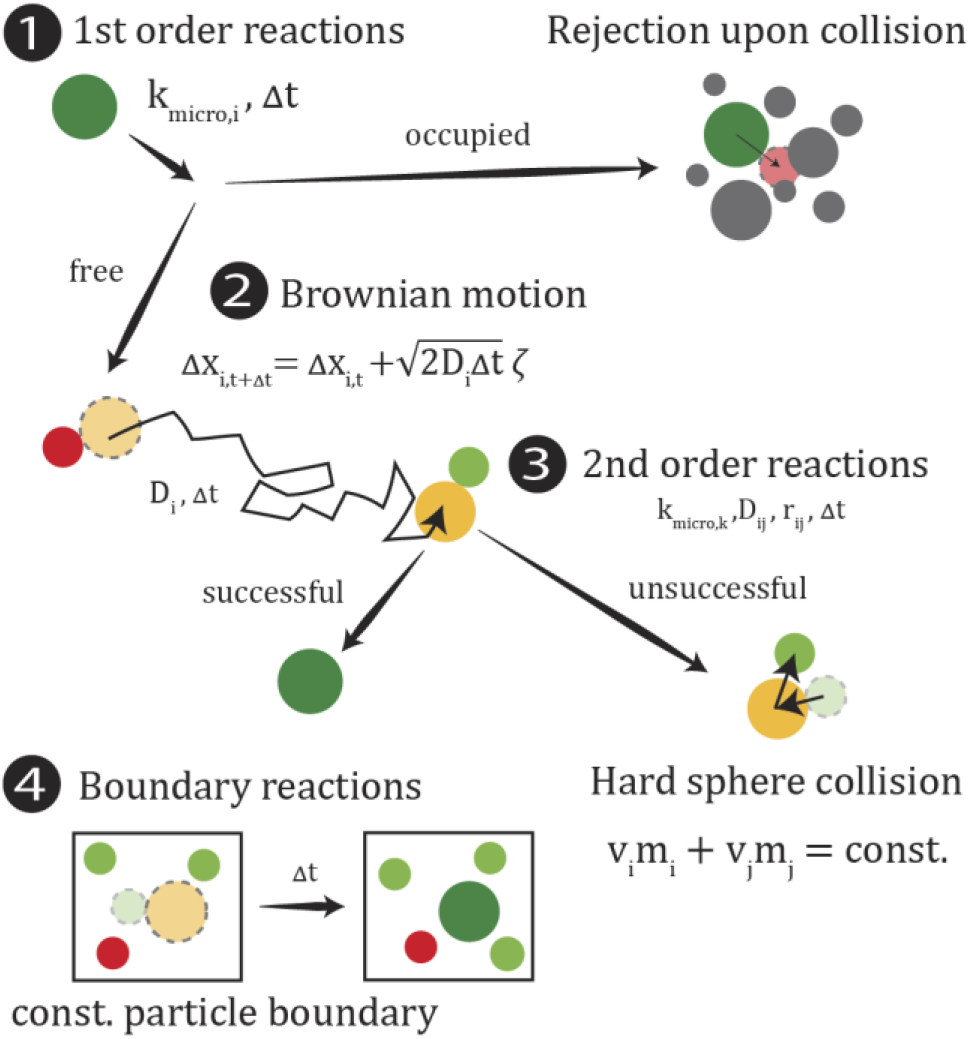
Individual algorithm steps of the molecular particle model. 1) 1^st^ order reactions are determined by a probabilistic success rate depending on the microscopic reaction rate per molecule *k_micro,i_* and the timestep Δ*t*. 2) The Brownian motion for every molecule is determined by its individual diffusion coefficient *D*_i_. 3) For 2^nd^ order reactions, success is determined upon collision given the microscopic reaction, the sum of the diffusion coefficient *D_ij_ = D_ij_ + D_j_*, the sum of the radii of the colliding particles *r_ij_* = *r_i_* + *r_j_*, and the time step (24). 4) Boundary reactions as constant particle boundaries, wherein the removal or insertion of particles is done if the number of particles deviates from the given boundary condition.

**Figure 2:**
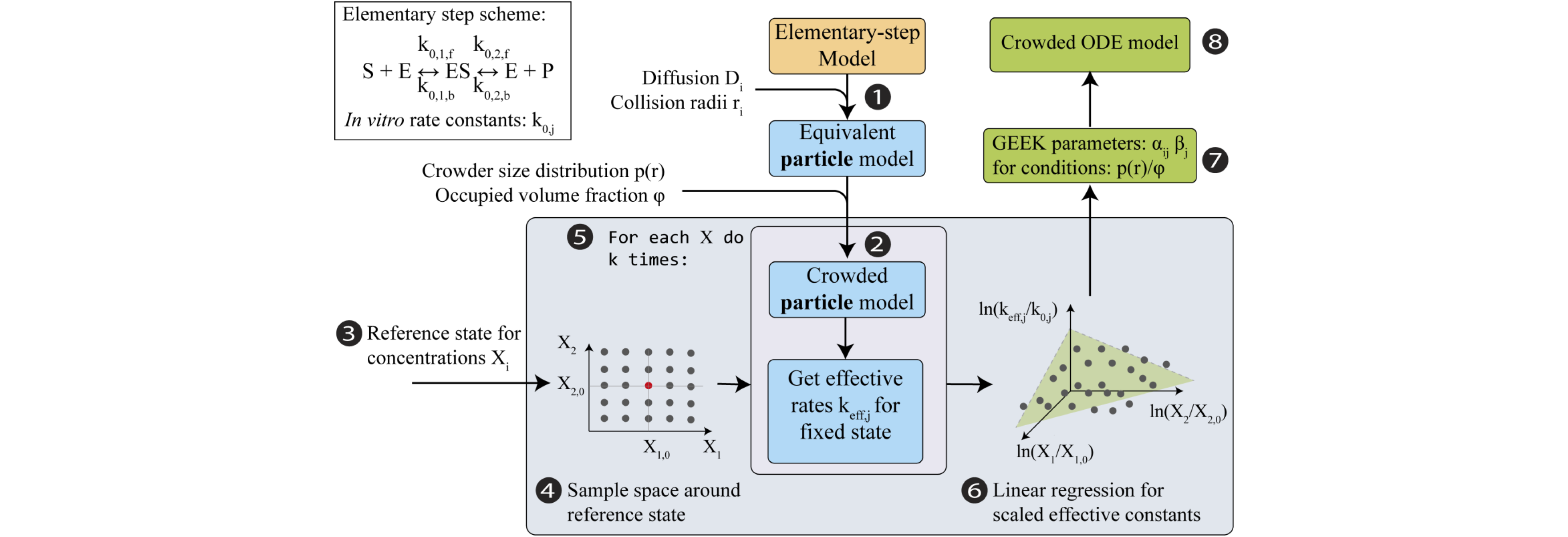
Modeling framework for crowded GEneralized Elementary Kinetics (GEEK). The input for the modeling framework is an arbitrary elementary step model containing *in vitro* data for the enzyme kinetics. 1) This model is then translated into an equivalent *in vitro* particle model of the enzymatic reaction. 2) The space is filled with inert molecules that are drawn from a size distribution *p*(*r*) until the fraction *ϕ* of the simulation space is occupied. 3) A reference concentration state is then chosen for the GEEK-model and 4) the space around the concentration space is sampled. 5) The *k* particle model realizations are then simulated for each concentration sample, i.e. repeat step 2 and simulate. 6) From the resulting particle traces, the effective rate constants are measured from the particle collision frequencies and the locally available volume. 7) These effective reaction rate constants are log-transformed, and a linear regression is performed with respect to the scaled logarithmic concentrations. The output of the linear regression directly links to the GEEK parameters, see Eq. (7) and Eq. (8). 8) Finally, the GEEK model can approximate the crowded kinetics using ordinary differential equations.

In the second step, the system is perturbed on the microscopic level to investigate the influence of crowding (Fig. 2, part 2). Therefore, inert particles are introduced into the system that therefore alter effective particle interactions between the reactive species (20). To model a realistic crowding environment, a size distribution function *p*(*r*) is estimated from the mass distribution *p*(*M_w_*) and an empirical mass size relation, 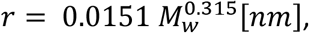, as reported for proteins in *E. coli* by Kalwarczyk *et al*. (36). The simulation volume is then populated with inert molecules by randomly drawing collision radii from the size distribution until the specified inert volume fraction *ϕ* is reached. The diffusion constant of the individual species is then calculated using the Stokes-Einstein relation, assuming that the hydrodynamic radius is equal to the collision radius:

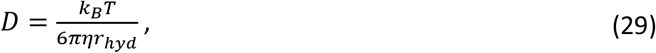

where *r_hyd_* is the hydrodynamic radius and η is the dynamic viscosity.

Next, the model is sampled around a chosen reference state (Fig. 2, parts 3 and 4). In this work, we chose to generate our sample with a full-factorial design. For each concentration sample, a particle simulation is performed where the effective rate constants *k_eff,j_* are measured for every elementary reaction as described in the previous section (Fig. 2, part 5).

Subsequently, multivariate linear regression is used to estimate the mean GEEK parameters *α_ij_* and *β_j_* for the specified crowding conditions (Fig. 2, part 6). Finally, the generalized elementary kinetics, as described above, are used to analyze the response behavior of an equivalent crowded ODE-enzyme model (Fig. 2, part 7).

### Weighted linear regressions

To estimate the GEEK parameters using multivariate regression, a multivariate regression was performed. Because the variance of the reaction rate would be expected to be dependent on the regression variables, a weighted linear regression was performed to avoid fitting data with a large heteroscedasticity (See Supplemental Text). The conditional variance of the residuals was therefore extracted, and a weighted linear regression was performed where each observation was weighted by the inverse of the conditional variance of the residual. To perform these calculations, the Python package *statsmodels* was used (37).

## Results & Discussion

To address the pitfalls currently associated with computational studies of enzymatic reactions in the intracellular space, this paper presents GEEK, a novel approach to capture spatial effects, such as crowding, in ordinary differential equation models. The framework is available in the form of two python packages: a package to implement GEEK expressions into ODE https://github.com/EPFL-LCSB/geek, and a package to perform openfpm-based hard-sphere Brownian reaction dynamics simulations https://github.com/EPFL-LCSB/openbread. The GEEK formulation directly quantifies the deviation from dilute mass-action behavior in a systematic and efficient procedure, and we have focused our studies on the impact of crowding due to the influence of densely packed biomolecules on enzyme reaction rates *in vivo*.

As an example, we applied our modeling framework to an investigation of the effects of macromolecular crowding on the enzymatic activity of PGM in *E. coli*. PGM is part of the lower glycolysis pathway and functions by reversibly transforming 3-phospho-D-glycerate (g3p) into 2-phospho-D-glycerate (g2p). We use PGM for our investigation as it exhibits a prototypical reversible Michaelis-Menten kinetics and it’s *in vitro* kinetics are well known (38).

### Impact of crowding on the elementary reaction level

For our reference elementary step mass-action model, which will serve as a basis for to construct the generalized elementary kinetic (GEEK) model, we calculated the elementary rate constants by the relations given in Eq. (3a,b) and Eq. (4a,b) from the *in vitro* Michaelis-Menten parameters measured by Fraser *et al*. (Table 1) (38). Based on this *in vitro* elementary step model, we built an equivalent *in vitro* particle model that required additional information on the molecular parameters, including mass, diffusion, and collision radius, of all the species involved in a reaction, meaning the substrates, products, free enzymes, and enzyme complexes. To estimate the collision radius of the enzyme and the enzyme-substrate complex, we followed the suggestions of Gameiro *et al*. and used the empirical relation between mass and size to estimate the inert molecule size from the enzyme mass (36, 39). In the same way, we applied the Stokes-Einstein relation to calculate the diffusion constants from the collision radius. We additionally assumed that the enzyme-substrate complex entirely enclosed the substrate with its binding pocket, thus rendering the collision radius of the complex and enzyme equal. To estimate the collision radius of g3p and g2p, we also followed the suggestions of Gameiro *et al*. and used the method developed by Zhao *et al*. to estimate their van der Waals volume and to calculate the equivalent sphere radius (39, 40). The diffusion constants of g2p and g3p were obtained from the literature (41). All molecular properties are summarized in Table 2.

**Table 1.** *In vitro* Michaelis-Menten parameters and calculated elementary rate constants for phosphoglycerate mutase in *E. coli*.

| Michaelis-Menten parameters |  | Elementary rate constants |  |
| --- | --- | --- | --- |
| $K_m$ g3p | 210 $\mu\text{M}$ (38) | $k_{1,f}$ | $15.2 \times 10^5 \text{ s}^{-1}\text{M}^{-1}$ |
| $K_m$ g2p | 97 $\mu\text{M}$ (38) | $k_{1,b}$ | $10 \text{ s}^{-1}$ |
| $k_{\text{cat}}$ g3p to g2p | $22 \text{ s}^{-1}$ (38) | $k_{2,f}$ | $22 \text{ s}^{-1}$ |
| $k_{\text{cat}}$ g2p to g3p | $10 \text{ s}^{-1}$ (38) | $k_{2,b}$ | $32.9 \times 10^5 \text{ s}^{-1}\text{M}^{-1}$ |

**Table 2.** Molecular properties of the reacting particles. † were calculated according to the approximations suggested by Gameiro *et al*. (39). The remaining values were obtained from (a) Perry (41) and Gameiro *et al*. (39) (b).

| Species | Diffusion ($\mu\text{m}^2\text{s}^{-1}$) | Collision radius (nm) | Mass (kDa) |
| --- | --- | --- | --- |
| g3p | 940 (a) | 1.11 † | 0.186 † |
| g2p | 940 (a) | 1.11 † | 0.186 † |
| PGM | 84.8 † | 3.87 † | 61 (b) |
| PGM complex | 84.8 † | 3.87 † | 61.186 |

Given the effective elementary rate constants and the molecular properties of the species, we calculated the effective microscopic rate constants using the relation given in Eq. (25) (Table 3). Comparing the microscopic rate constants in Table 3 with the diffusion-limited rate constants, it can be seen that the diffusion limited constants γ_1,*f*_ = γ_2,*b*_ = 3.88 × 10^10^ M^−1^*S*^−1^ are about five orders of magnitude higher than the microscopic reaction constants. This indicates that the microscopic binding process is much slower than the diffusion process and that the kinetics are reaction limited and not diffusion limited. Thus the mean time until the first collision between two reactants, i.e. the mean first passage time, is orders of magnitudes shorter than the mean time to the first reaction. For a reaction to be successful tens of thousands of collisions are occurring, hence, the impact of any increase in high-frequency first passage events due to fractal diffusion is limited (22). As most enzymes are reaction limited, the effects originating from fractal diffusion are therefore not likely dominating in metabolic networks.

**Table 3.** Microscopic reaction rates per reacting particle (p), per collision (c), and diffusion-limited rate constants of the bimolecular reaction

| Microscopic rate constants |  |
| --- | --- |
| $k_{1,f}$ (c) | $15.7 \times 10^5 \text{ s}^{-1}\text{M}^{-1}$ |
| $k_{1,b}$ (p) | $10 \text{ s}^{-1}$ |
| $k_{2,f}$ (p) | $22 \text{ s}^{-1}$ |
| $k_{2,b}$ (c) | $34.0 \times 10^5 \text{ s}^{-1}\text{M}^{-1}$ |

To build a GEEK model, that allows to characterize the enzyme kinetics in a crowded environment, we sampled the concentration space. This was done using a full-factorial design, allowing us to study the effect of several variables on the response output, as well as interactions between those variables, that sampled both the product and substrate concentrations as well as different enzyme saturation levels, indicating the percentage of bound enzyme with respect to the total enzyme concentration. For the regression input space, all combinations of substrate and product concentrations that were n-fold increased and decreased with respect to the reference concentration of [*S*]_0_ = [*P*]_0_ = 49 μM were used, with *n* ∊ [1,2,3,4,5], in combination with all free enzyme and enzyme-complex concentrations that yielded saturations of *σ* = [0.1,0.2,0.3,0.4,0.5,0.6,0.7,0.8,0.9] given a total amount of enzyme [*E_tot_*] = 64 μM. Furthermore, ten independent realizations of the crowding population were used for every sample concentration to capture the variability that comes from differently sized crowding-agents drawn from the size distribution. Thus, we measured the effective rate constants ten times for each of the 729 concentration samples and crowding conditions.

In total, we generated 21 generalized elementary kinetic models for five different inert volume fractions *ϕ_k_* and four different size distributions *p_k_*(*r*) plus one without any crowding. This allowed for a detailed comparison of the effects of the volume fraction and size distribution of the crowding agents on enzyme kinetics. For the size-distributions, we used (i) the *E. coli* distribution derived from Kalwarczyk *et al*., (ii) a population containing only particles of the median size of the *E. coli* distribution, (iii) a population the size of the upper quartile of the *E. coli* size distribution, and (iv) a population the size of the lower quartile (Fig. 3) (36). These crowding populations were each investigated for inert volume fractions of *ϕ* ∊ [0.0,0.1,0.2,0.3,0.4,0.5].

**Figure 3:**
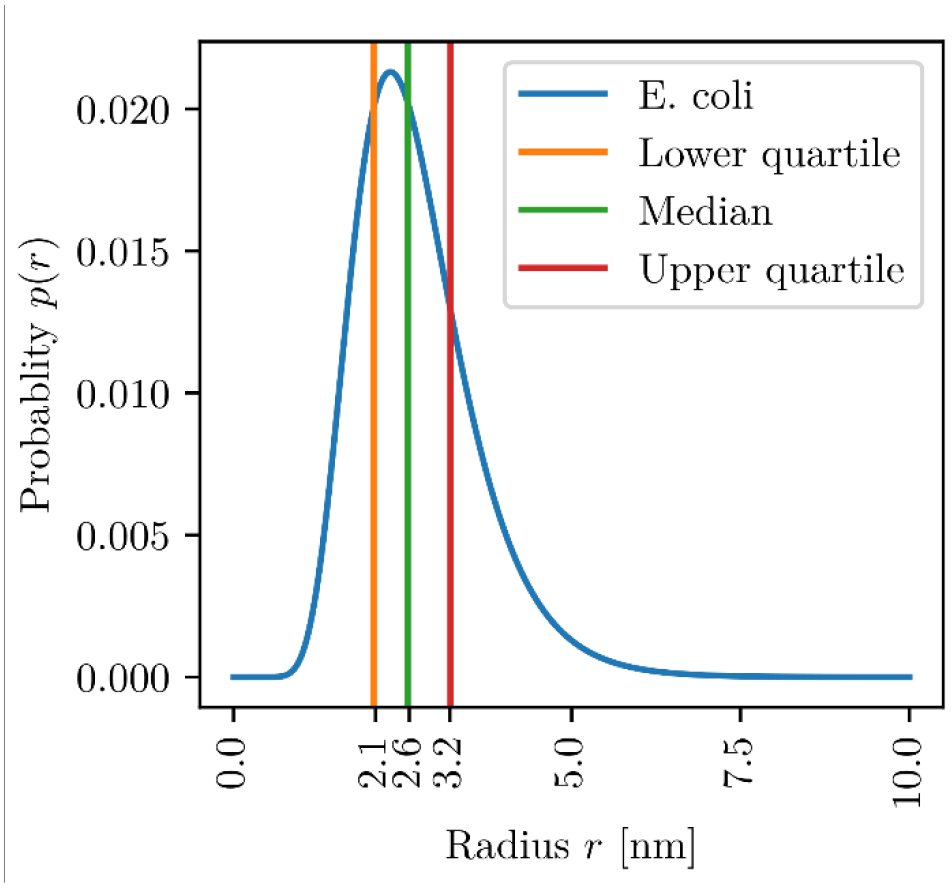
Size distribution function of the inert particles, numerically calculated from the mass distribution and empirical mass size relation as reported for proteins in *E. coli* by Kalwarczyk *et al*. (36).

For each crowding condition, we estimated the mean GEEK parameters *α_ij_ and β_j_* using multivariate weighted linear regression, which indicate conditions that likely influenced the enzyme kinetics. To further quantify the uncertainty of the mean GEEK parameters, the 95% confidence intervals of the regression results are given in Table 4. For parameter estimates with a p value ≥ 0.05, it was assumed that no significant correlation existed, and these parameters were not accounted for in the GEEK model. Note that this mean GEEK parameter model assumes that the crowding composition of an average cell is given by the average effect of a crowding configuration on the rate constant, which should be accurate for cell populations of the because the average has been shown to be reflective of the overall state of a cell population.

**Table 4.** Parameters of the generalized elementary kinetics, i.e. *α_ij_* and *β_j_* and, for all elementary reactions at different inert volume fractions. The dashes denote GEEK parameters with a significance of *p* ≤ 0.05.

| Elementary reaction |  | Crowding conditions |  |  |  |  |  |
| --- | --- | --- | --- | --- | --- | --- | --- |
| Formula | Parameters | 0% | 10% | 20% | 30% | 40% | 50% |
| $S + E \xrightarrow{k_{1,f}} ES$ | $\beta_{1,f}$ | $7.31 \times 10^{-2}$ | $2.61 \times 10^{-1}$ | $4.57 \times 10^{-1}$ | $6.87 \times 10^{-1}$ | $9.57 \times 10^{-1}$ | 1.25 |
| | $\alpha_{S,1,f}$ | $4.74 \times 10^{-3}$ | — | $3.22 \times 10^{-3}$ | $7.22 \times 10^{-3}$ | $6.95 \times 10^{-3}$ | $3.23 \times 10^{-3}$ |
| | $\alpha_{E,1,f}$ | $1.07 \times 10^{-2}$ | $1.89 \times 10^{-2}$ | $9.67 \times 10^{-3}$ | $8.07 \times 10^{-3}$ | $1.23 \times 10^{-2}$ | — |
| | $\alpha_{ES,1,f}$ | — | $1.34 \times 10^{-2}$ | — | — | — | — |
| | $\alpha_{P,1,f}$ | — | $2.51 \times 10^{-3}$ | $-3.47 \times 10^{-3}$ | — | — | $2.42 \times 10^{-3}$ |
| $ES \xrightarrow{k_{1,b}} S + E$ | $\beta_{1,b}$ | $-1.48 \times 10^{-2}$ | $-1.17 \times 10^{-1}$ | $-2.80 \times 10^{-1}$ | $-5.46 \times 10^{-1}$ | -1.03 | -2.26 |
| | $\alpha_{S,1,b}$ | $-2.94 \times 10^{-3}$ | $-3.56 \times 10^{-3}$ | $-4.53 \times 10^{-3}$ | $-6.09 \times 10^{-3}$ | $-8.33 \times 10^{-3}$ | $-9.58 \times 10^{-3}$ |
| | $\alpha_{E,1,b}$ | $-1.90 \times 10^{-4}$ | $1.71 \times 10^{-3}$ | $6.47 \times 10^{-4}$ | $3.32 \times 10^{-3}$ | — | $3.16 \times 10^{-3}$ |
| | $\alpha_{ES,1,b}$ | — | $3.46 \times 10^{-3}$ | — | $2.03 \times 10^{-3}$ | $-1.22 \times 10^{-3}$ | — |
| | $\alpha_{P,1,b}$ | $-2.94 \times 10^{-3}$ | $-3.58 \times 10^{-3}$ | $-4.52 \times 10^{-3}$ | $-6.09 \times 10^{-3}$ | $-8.33 \times 10^{-3}$ | $-9.58 \times 10^{-3}$ |
| $ES \xrightarrow{k_{2,f}} P + E$ | $\beta_{2,f}$ | $-1.48 \times 10^{-2}$ | $-1.17 \times 10^{-1}$ | $-2.80 \times 10^{-1}$ | $-5.46 \times 10^{-1}$ | -1.03 | -2.26 |
| | $\alpha_{S,2,f}$ | $-2.94 \times 10^{-3}$ | $-3.56 \times 10^{-3}$ | $-4.53 \times 10^{-3}$ | $-6.09 \times 10^{-3}$ | $-8.34 \times 10^{-3}$ | $-9.58 \times 10^{-3}$ |
| | $\alpha_{E,2,f}$ | $-1.90 \times 10^{-4}$ | $1.71 \times 10^{-3}$ | $6.53 \times 10^{-4}$ | $3.31 \times 10^{-3}$ | — | $3.13 \times 10^{-3}$ |
| | $\alpha_{ES,2,f}$ | — | $3.46 \times 10^{-3}$ | — | $2.01 \times 10^{-3}$ | $-1.20 \times 10^{-3}$ | — |
| | $\alpha_{P,2,f}$ | $-2.94 \times 10^{-3}$ | $-3.58 \times 10^{-3}$ | $-4.52 \times 10^{-3}$ | $-6.10 \times 10^{-3}$ | $-8.34 \times 10^{-3}$ | $-9.57 \times 10^{-3}$ |
| $P + E \xrightarrow{k_{2,b}} ES$ | $\beta_{2,b}$ | $6.88 \times 10^{-2}$ | $2.54 \times 10^{-1}$ | $4.58 \times 10^{-1}$ | $6.85 \times 10^{-1}$ | $9.52 \times 10^{-1}$ | 1.25 |
| | $\alpha_{S,2,b}$ | — | — | $-2.29 \times 10^{-3}$ | — | $3.09 \times 10^{-3}$ | — |
| | $\alpha_{E,2,b}$ | — | — | $1.05 \times 10^{-2}$ | — | — | — |
| | $\alpha_{ES,2,b}$ | — | — | — | $-7.78 \times 10^{-3}$ | — | — |
| | $\alpha_{P,2,b}$ | $6.59 \times 10^{-3}$ | $2.76 \times 10^{-3}$ | $4.70 \times 10^{-3}$ | $6.85 \times 10^{-3}$ | $8.45 \times 10^{-3}$ | $3.74 \times 10^{-3}$ |

Comparing the parameters *α_ij_* and *β_j_* of each elementary reaction *j* (Table 4), it can generally be observed that the direct effect *β_j_* is about one to two orders of magnitude larger than each corresponding coupling coefficient, *α_ij_*. Therefore, the direct effect is on the order of ±10^−2^ to ±10^0^, whereas the coupling coefficients are on the order of ±10^−4^ to ±10^−2^. Assuming a two-fold increase in a concentration, the change in the coupling is smaller than one percent, whereas the direct effect varies between one and a thousand percent. This suggests that the effect of the reduced dimensionality only plays a small role compared to the effective interaction potential and the diffusion inhibition. For comparison, we show in the supplementary material that for diffusion-limited reactions, GEEK captures the influence of fractal diffusion as a deviation in the effective order as described by the coupling coefficient α_i,j_ (Supplementary Fig. 5) (16, 22, 42).

### Effects of crowding on the reversible Michalis-Menten kinetics

We used the results of the linear regression to parameterize GEEK models to compare the ODE-simulations of the classical Michaelis-Menten experiment with the mass-action model. The basis of this experiment involved an initial substrate concentration [*S*]_*init*_ that was added to a volume with a fixed enzyme concentration [*E*]_*init*_ = [*E*]_*tot*_. When the substrate was added, the enzyme started to convert the substrate into a product. If the enzyme was operating reversibly, part of the product would also be converted back to a substrate, and the reaction would become indistinguishable as it approached equilibrium. In this equilibrium state, the overall free energy of the reaction *Δ_r_G°’* was close to zero. Therefore, the ratio between the product and substrate concentrations could be used to estimate the apparent equilibrium constant *K_eq_*.

To characterize the dynamics of this system, the time to half-equilibrium *t_eq_*_/2_ was measured, which indicates the time needed for the ratio between the product and substrate concentrations to equal *K_eq_*/2 (Fig. 4a). In general, an increase in the *t_eq_*_/2_ was seen with an increasing substrate concentration (Fig. 4a). The time to half-equilibrium for the interconversion between g3p to g2p was reduced for small substrate concentrations and inert molecule fractions, to up *ϕ* = 30 –40%. In the case of [*S_init_* = [*S*]*_ref_*/4, the time to half-equilibrium was reduced to a minimum value for an inert volume fraction of *ϕ* = 40% (Fig. 4a). For [*S*]_init_ = [*S*]_*ref*_ and [*S*]_init_ = 2[*S*]_*ref*_ this decrease in half-life time persists, though the overall half-life times are larger than for [*S*]_init_ = [*S*]_*ref*_/4 and the minimum point occurs at lower inert volume fractions. Finally, in the [S]_init_ = 4[*S*]*_ref_* case, this decrease in *t_eq_*_/2_ is no longer visible. It follows from this that the average initial rate increases with substrate concentration and decreases with an increasing volume occupancy. This suggest that the same substrate concentrations yield higher enzyme saturations, meaning that the ratio of enzyme-substrate complex to the total amount of enzyme increases and that the dissociation of the enzyme-substrate complex is inhibited.

**Figure 4:**
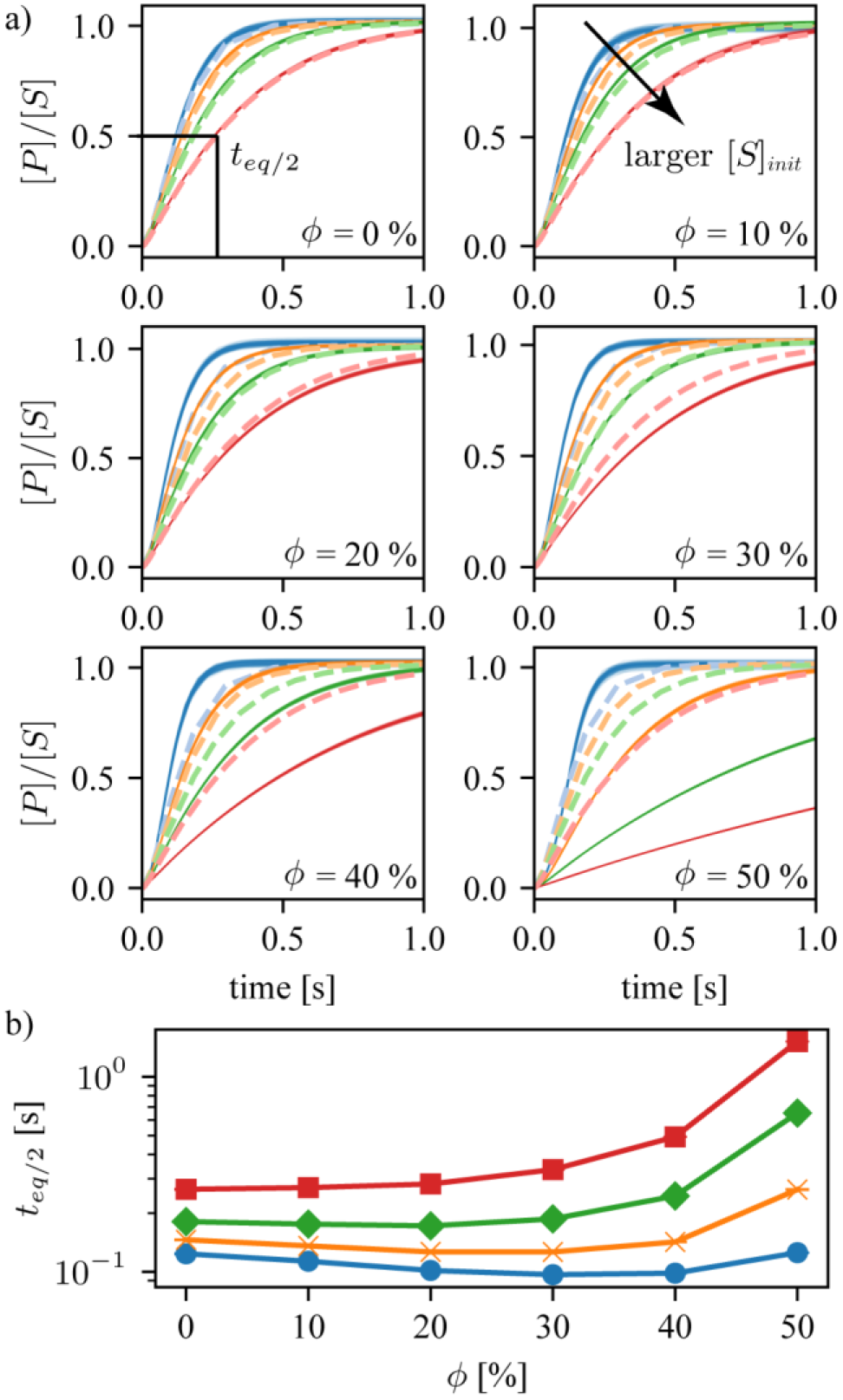
a)[*P*]/[*S*] Dynamics determined for mass-action and generalized elementary kinetics (GEEK) models for different initial substrate concentrations [*S*]_init_ and different occupied volume fractions (*ϕ*)for the *E. coli* molecular weight distributions. The light dashed lines represent the dilute mass-action model, whereas the thin solid lines represent a population of 100 re-sampled GEEK models. b) Time to half-equilibrium *t_eq_*_/2_ as a function of the occupied volume fraction for different initial substrate concentrations [*S*]_init_. The colors of the lines denote the different initial concentrations, where blue corresponds to [*S*]_*init*_ = [*S*]_*ref*_/4, yellow to [*S*]_*init*_ = [*S*]_*ref*_, green to [*S*]_*init*_ = 2[*S*]_*ref*_, and red to [*S*]_*init*_ = 4[*S*]_ref_.

**Figure 5:**
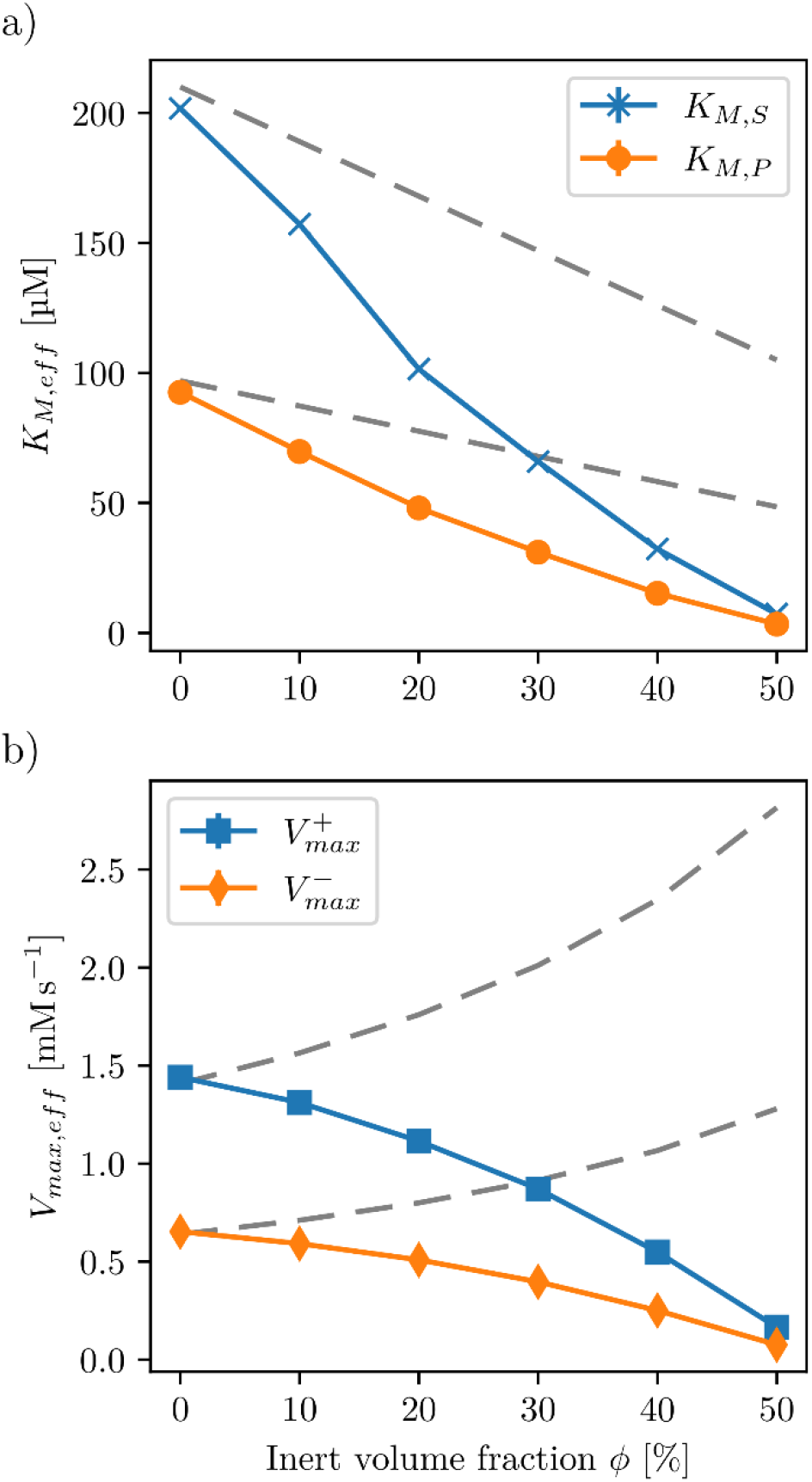
Effective Michaelis-Menten parameters a) *K_M,S_* and *K_M,P_* and 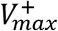 and 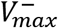 as a function of volume fraction. The grey dashed lines represent the effective parameters when the all concentrations are scaled to an effective volume *V_eff_ = V*(1 — *ϕ*) that excludes the volume occupied by inert molecules. The errors in the values calculated from uniformly resampling the GEEK parameters are smaller than 2% of the mean.

**Figure 6:**
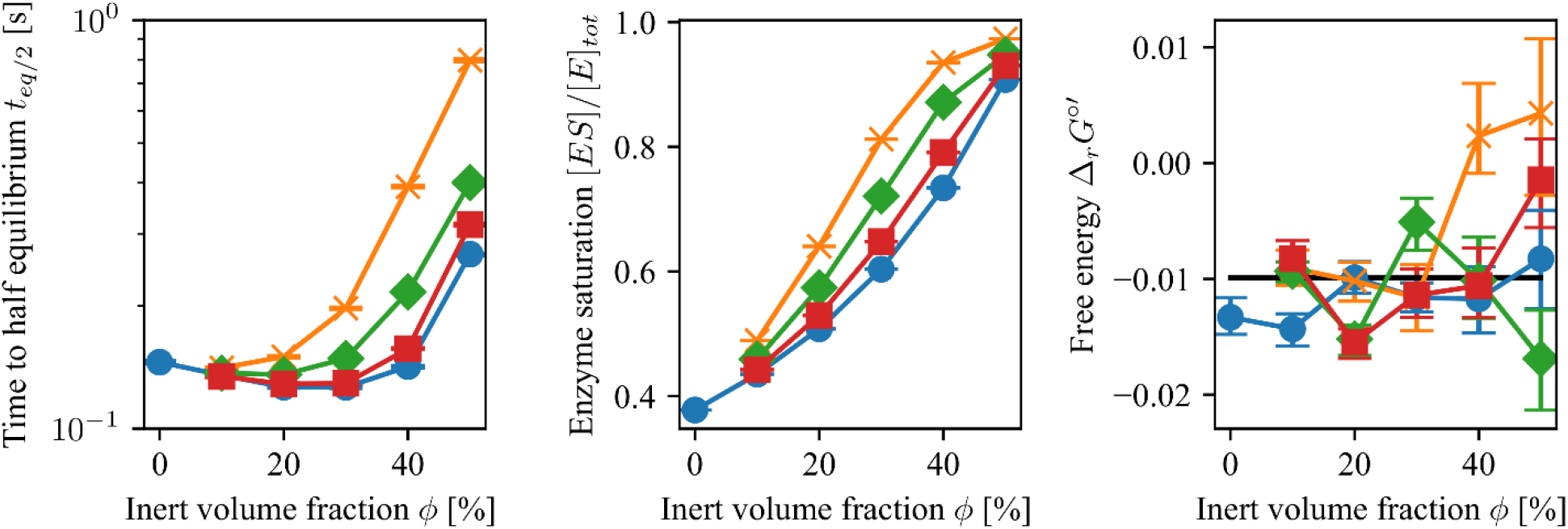
Effects of the particle size distribution. a) Time to half-equilibrium *t_eq_*_/2_, b) enzyme saturation [*ES*]/[*E*]_*tot*_ for [*S*]_*ref*_ = 49 μM, and c) apparent standard free energy of the reaction measured as *RT*log([*P*]_*eq*_/[*S*]_*eq*_) under different crowding conditions. The blue line represents the apparent equilibrium measured from the *E. coli* size distribution; yellow, green, and red are obtained using a single size of inert molecules corresponding to the lower quartile, the median quartile, and the upper quartile of the *E. coli* size distribution, respectively. The error bars denote the upper and lower quartile of the resulting population that was obtained by resampling the GEEK model parameters within their confidence bounds. The black line denotes the equilibrium constant calculated from the *in vitro* kinetic parameters.

**Figure 7:**
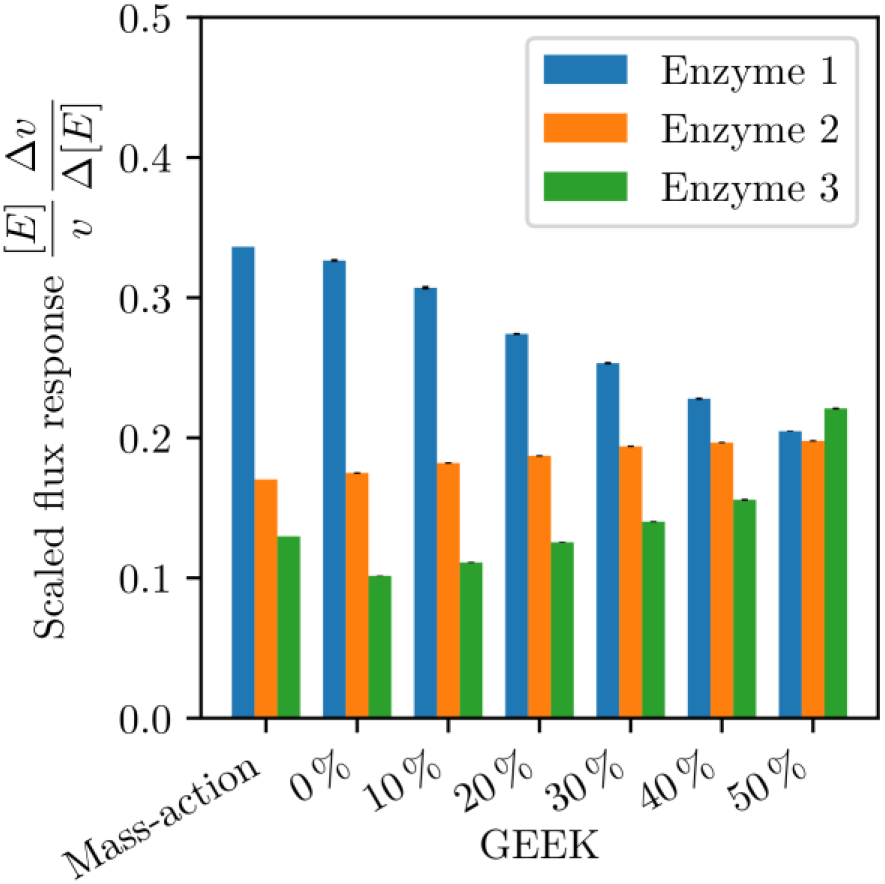
Flux responses corresponding to a two-fold increase in the respective enzyme concentration for the basic mass-action model and the GEEK models derived from the *E. coli* distributions as well as the different volume fractions of the inert molecules. Blue denotes the first enzyme in the pathway, orange the second, and green the third.

For a closer analysis of these findings, the Michaelis-Menten parameters were estimated using Eadie-Hofstee diagrams, solving for the steady-state flux of the substrate and product concentrations between 4.9 μM and 490 μM, (Eq.(11)). The Eadie-Hofstee diagrams (Fig. 5) reveal that for both high and low occupancy volume fractions, a slight non-linearity with respect to the linear Eadie-Hofstee form of the reversible Michaelis-Menten is introduced with the generalized elementary kinetics. This indicates that the effective maximal flux 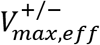 and effective Michealis-Menten constant *K_M,X,eff_* are actually functions of the reactant concentrations [*S*] and [*P*]. For the case of *ϕ* = 0%, this non-linearity is only pronounced at small reactant concentrations, whereas for higher volume occupancy conditions, the non-linearty is visible over the entire measurement range. Nevertheless, we used linear regression to estimate the effective average parameters to compare the steady-state GEEK model to the traditional Michaelis-Menten kinetics (Supplementary Fig. 3).

Interestingly, the steady-state analysis revealed that all the effective Michaelis-Menten parameters 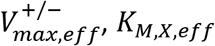, *K_M,X,eff_* decreased as a function of the inert occupied volume *ϕ* These results complement our primary analysis, as the maximal flux of the enzyme directly relates to the ability of the enzyme-substrate complex to dissociate, and the Michaelis-Menten constant is a measure of the affinity of the reactant binding to the enzyme. The lower the Michaelis-Menten constant, the higher the binding affinity to the enzyme. Consequently, a decreasing Michaelis-Menten constant indicates more enzyme bound at the same reactant concentration, or in other words, an increased enzyme saturation. For the effective flux through the enzyme, this results in two counteracting effects: a potential increase in flux due to an increase in saturation or a decrease in flux due to the reduced dissociation. From the analysis of t_eq/2_ and the effective Michaelis-Menten parameters, it is evident that the flux-increasing effect is dominating if enough free enzyme is available to increase the saturation. If the enzyme capacity does not allow more substrate to associate, the flux-decreasing effect dominates.

### Influence of crowder size on the Michaelis-Menten kinetics

We further investigated the influence of the size of the inert molecules on the enzyme kinetics by comparing crowding with different inert molecule sizes obtained using the results for the *E. coli* size distribution. When comparing the *t_eq_*_/2_ as a function of the volume occupancy obtained from crowding using the *E. coli* distribution to the population consisting of a single inert molecule size, a general flux-decreasing effect was observed for crowding in the single-sized population. Furthermore, smaller inert molecule sizes showed a stronger flux-decreasing effect that was alleviated as the size of the inert molecules increased. When we compared the enzyme saturation at equilibrium for the different inert molecule sizes, the single-sized crowding showed an increased saturation, and smaller crowding sizes had a stronger effect. This shows that the overall substrate affinity is increased more if the inert molecules are smaller than the enzyme-substrate collision radius.

Finally, we determined the effective standard free energy of the reaction from the effective equilibrium constant Δ*_r_G°’* = – *RT*ln(*K_eq_*), where the effective equilibrium constant was determined from the reactant concentrations at equilibrium *K_eq_* = [*P*]*_eq_*/[*S*]*_eq_*. This showed that the overall apparent standard free energy of reaction does not vary significantly with crowding size or volume fraction. Since the non-ideal contributions, which contain the terms emerging from molecular interactions, from steric interaction for the substrates and products are exactly equal, we would expect the overall non-ideal contribution to the free energy of the enzymatic reaction to be zero. The deviation in the effective standard energy using GEEK can be attributed to the approximation over the state space at points far from equilibrium (See Supplementary Fig. 7).

### Consequences of the crowded kinetics on the pathway level

We further investigated the effects of crowding at a system level using an example linear pathway with three enzymes. We considered a linear pathway where a compound *X_n_* was reversibly converted into a compound *X_n_*_+1_. All reactions were considered to follow reversible Michaelis-Menten kinetics.

The concentration of the first compound [*X*_1_] was considered to be 250 μM and the concentration of the last compound [*X*_4_] to be 50 μM. These boundary concentrations were considered to be constant. The last reaction catalyzed by enzyme 3 was parameterized using the results found for PGM. We further choose the parameters of enzyme 1 and 2 by scaling the product-specific Michaelis-Menten constant *K_M,P_* by a factor of 2 and 3, respectively, as well as the maximum backwards flux 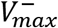 by a factor of 0.5 and 0.33, respectively (Table 5). We further considered that all enzymes and reactants were of the same size and that, therefore, their GEEK parameters *α_ij_* and *β_j_* were assumed to be the same. This models short pathway where only minor modifications occur on the molecule for example the reallocation of a phosphate group or a double bond.

**Table 5.** Enzyme parametrizations used for the linear pathway example.

| Parameters | Enzyme 1 | Enzyme 2 | Enzyme 3 |
| --- | --- | --- | --- |
| $K_{M,S}$ | 200 μM | 200 μM | 200 μM |
| $K_{M,P}$ | 300 μM | 200 μM | 100 μM |
| $V_{max}^+$ | 1.5 mM/s | 1.5 mM/s | 1.5 mM/s |
| $V_{max}^-$ | 0.15 mM/s | 0.25 mM/s | 0.5 mM/s |

To characterize the influence of crowding on our pathway, we calculated the relative responses of the flux with respect to a two-fold increase in each of the enzyme concentrations, modeling an overexpression of the respective enzyme. We then compared the results obtained for the traditional mass-action model with the GEEK models representing the *E. coli* size distribution at different occupied volume fractions. Comparing the GEEK model without any inert molecules with the results from the mass-action model, only minor differences were observed. On the other hand, a redistribution of the flux control was clearly seen for increasing volume fractions, wherein the initially largest flux response decreased and the lower flux responses increased as a function of the volume fraction. Considering the typical volume occupancy in cells of 20% to 40%, the relative order between the flux responses was still the same as under dilute conditions, though the magnitude of the largest flux response is significantly reduced. In this case, it is the relative sensitivity of the flux that is reduced. This could act as an additional stabilization mechanism with respect to fluctuations in the enzyme levels, though it should be noted that this effect does depend on many factors, such as relative *K_M_* s, *V_max_*, and reaction *Δ_r_G’s*. How these factors impact the response of the pathway fluxes with respect to the change in enzyme levels will require an extensive analysis considering different pathway structures and different types of enzyme kinetics. Given the extend of such a study it will beneficial to first understand impact for different types of enzyme kinetics before moving to the network level.

## Conclusion

This research presents a method for characterizing spatial effects of any nature on biochemical reactions based on the mapping of average effects to ordinary differential equations. Besides studying the effects of intracellular crowding as we have done, this framework can influence the study of membrane confined biochemical reactions, enzyme-channeling, and DNA or actin bound reactions systems, which are all current topics in biochemistry that lack dedicated study tools. Using a representative example, we confirm the hypothesis of recent research in the field that for reaction-limited enzyme kinetics, the diffusion effects in fractal spaces are negligible and are most likely not dominating in reaction networks. Instead, a strong direct effect of crowding was seen on the effective rates, where a decrease in dissociation rates and an increase in association rates was observed when increasing occupied volume fraction. Both effects can be sufficiently explained by an effective increase of the crowding induced potential with the volume fraction, confirming that this is a better predictor of intracellular enzyme kinetics than the diffusion. Furthermore, we show that the effective Michaelis-Menten parameters strongly depended on the volume occupancy and the size distribution of inert molecules, indicating that the kinetics are likely to vary dramatically in different cellular compartments. We finally show that crowding at a simplified network level can lead to a redistribution of the effective control on the flux response, suggesting that crowding can have a stabilizing effect with respect to fluctuation in enzyme levels, potentially indicating why enzymatic systems *in vivo* systems show a higher robustness compared to *in vitro*.

In future work, this framework will be used to analyze the impact of crowding on other kinetic mechanisms and on an expanded network level. The results will illuminate the strength of the overall impact of crowding on the regulation of metabolism.

## Authors contributions

D.W. designed the model and performed the simulations, V.H. conceived the original idea and supervised the project. Both D.W and V.H. authors contributed to the final version of the manuscript.

## Acknowledgements

This project has received funding from the European Union’s Horizon 2020 research and innovation program under grant agreement No 686070. The authors would like to thank the MOSAIK Group (Center for Systems Biology Dresden) in particular Pietro Incardona for technical support on OPENFPM

## References

1. Ellis, R. J. 2001. Macromolecular crowding: obvious but underappreciated. Trends in Biochemical Sciences 26(10):597–604.

2. Minton, A. P. 2001. The influence of macromolecular crowding and macromolecular confinement on biochemical reactions in physiological media. J Biol Chem 276(14):10577–10580.

3. Zhou, H. X., and S. Qin. 2013. Simulation and Modeling of Crowding Effects on the Thermodynamic and Kinetic Properties of Proteins with Atomic Details. Biophys Rev 5(2):207–215.

4. Aon, M. A., and S. Cortassa. 2015. Function of metabolic and organelle networks in crowded and organized media. Frontiers in Physiology 5.

5. Hancock, R. 2004. Internal organisation of the nucleus: assembly of compartments by macromolecular crowding and the nuclear matrix model. Biol Cell 96(8):595–601.

6. Poggi, C. G., and K. M. Slade. 2015. Macromolecular crowding and the steady-state kinetics of malate dehydrogenase. Biochemistry-Us 54(2):260–267.

7. Yadav, J. K. 2013. Macromolecular Crowding Enhances Catalytic Efficiency and Stability of alpha-Amylase. ISRN Biotechnol 2013:737805.

8. van den Berg, B., R. J. Ellis, and C. M. Dobson. 1999. Effects of macromolecular crowding on protein folding and aggregation. EMBO J 18(24):6927–6933.

9. Emiola, A., J. George, and S. S. Andrews. 2015. A Complete Pathway Model for Lipid A Biosynthesis in Escherichia coli. Plos One 10(4).

10. Watterson, S., M. L. Guerriero, M. Blanc, A. Mazein, L. Loewe, K. A. Robertson, H. Gibbs, G. H. Shui, M. R. Wenk, J. Hillston, and P. Ghazal. 2013. A model of flux regulation in the cholesterol biosynthesis pathway: Immune mediated graduated flux reduction versus statin-like led stepped flux reduction. Biochimie 95(3):613–621.

11. Andreozzi, S., A. Chakrabarti, K. C. Soh, A. Burgard, T. H. Yang, S. Van Dien, L. Miskovic, and V. Hatzimanikatis. 2016. Identification of metabolic engineering targets for the enhancement of 1,4-butanediol production in recombinant E. coli using large-scale kinetic models. Metab Eng 35:148–159.

12. Andreozzi, S., L. Miskovic, and V. Hatzimanikatis. 2016. iSCHRUNK – In Silico Approach to Characterization and Reduction of Uncertainty in the Kinetic Models of Genome-scale Metabolic Networks. Metab Eng 33:158–168.

13. Khodayari, A., A. R. Zomorrodi, J. C. Liao, and C. D. Maranas. 2014. A kinetic model of Escherichia coli core metabolism satisfying multiple sets of mutant flux data. Metab Eng 25:50–62.

14. Brooks, H. B., S. Geeganage, S. D. Kahl, C. Montrose, S. Sittampalam, M. C. Smith, and J. R. Weidner. 2004. Basics of Enzymatic Assays for HTS. Assay Guidance Manual. G. S. Sittampalam, N. P. Coussens, K. Brimacombe, A. Grossman, M. Arkin, D. Auld, C. Austin, J. Baell, B. Bejcek, T. D. Y. Chung, J. L. Dahlin, V. Devanaryan, T. L. Foley, M. Glicksman, M. D. Hall, J. V. Hass, J. Inglese, P. W. Iversen, S. D. Kahl, S. C. Kales, M. Lal-Nag, Z. Li, J. McGee, O. McManus, T. Riss, O. J. Trask, Jr., J. R. Weidner, M. Xia, and X. Xu, editors, Bethesda (MD).

15. Zimmerman, S. B., and S. O. Trach. 1991. Estimation of Macromolecule Concentrations and Excluded Volume Effects for the Cytoplasm of Escherichia-Coli. J Mol Biol 222(3):599–620.

16. Schnell, S., and T. E. Turner. 2004. Reaction kinetics in intracellular environments with macromolecular crowding: simulations and rate laws. Progress in Biophysics & Molecular Biology 85(2–3):235–260.

17. Grima, R., and S. Schnell. 2006. A systematic investigation of the rate laws valid in intracellular environments. Biophys Chem 124(1):1–10.

18. Klann, M. T., A. Lapin, and M. Reuss. 2011. Agent-based simulation of reactions in the crowded and structured intracellular environment: Influence of mobility and location of the reactants. Bmc Systems Biology 5.

19. Mourao, M., D. Kreitman, and S. Schnell. 2014. Unravelling the impact of obstacles in diffusion and kinetics of an enzyme catalysed reaction. Physical chemistry chemical physics: PCCP 16(10):4492–4503.

20. Berezhkovskii, A. M., and A. Szabo. 2016. Theory of Crowding Effects on Bimolecular Reaction Rates. J Phys Chem B 120(26):5998–6002.

21. Galanti, M., D. Fanelli, S. D. Traytak, and F. Piazza. 2016. Theory of diffusion-influenced reactions in complex geometries. Physical Chemistry Chemical Physics 18(23):15950–15954.

22. Benichou, O., C. Chevalier, J. Klafter, B. Meyer, and R. Voituriez. 2010. Geometry-controlled kinetics. Nat Chem 2(6):472–477.

23. Shim, A. R., L. Almassalha, H. Matsuda, R. Nap, and I. Szleifer. 2017. Dynamic modeling shows long-term gene expression is highly dependent on macromolecular crowding. Faseb J 31.

24. Bar-Even, A., E. Noor, Y. Savir, W. Liebermeister, D. Davidi, D. S. Tawfik, and R. Milo. 2011. The moderately efficient enzyme: evolutionary and physicochemical trends shaping enzyme parameters. Biochemistry-Us 50(21):4402–4410.

25. Feig, M., and Y. Sugita. 2013. Reaching new levels of realism in modeling biological macromolecules in cellular environments. J Mol Graph Model 45:144–156.

26. Heinrich, R., and S. Schuster. 1996. The regulation of cellular systems. Chapman & Hall, New York; London.

27. Eadie, G. S. 1942. The inhibition of cholinesterase by physostigmine and prostigmine. J Biol Chem 146(1):85–93.

28. Hofstee, B. H. J. 1959. Non-Inverted Versus Inverted Plots in Enzyme Kinetics. Nature 184(4695):1296–1298.

29. Demirel, Y. 2014. Nonequilibrium thermodynamics: transport and rate processes in physical, chemical and biological systems. Elsevier, Amsterdam.

30. Strating, P. 1999. Brownian dynamics simulation of a hard-sphere suspension. Phys Rev E 59(2):2175–2187.

31. Allen, M. P. 1980. Brownian Dynamics Simulation of a Chemical-Reaction in Solution. Mol Phys 40(5):1073–1087.

32. Northrup, S. H., S. A. Allison, and J. A. Mccammon. 1984. Brownian Dynamics Simulation of Diffusion-Influenced Bimolecular Reactions. Journal of Chemical Physics 80(4):1517–1526.

33. Wang, M. C., and G. E. Uhlenbeck. 1945.On the Theory of the Brownian Motion-Ii. Rev Mod Phys 17(2–3):323–342.

34. Morelli, M. J., and P. R. ten Wolde. 2008. Reaction Brownian dynamics and the effect of spatial fluctuations on the gain of a push-pull network. Journal of Chemical Physics 129(5).

35. Collins, F. C., and G. E. Kimball. 1949. Diffusion-Controlled Reactions in Liquid Solutions. Ind Eng Chem 41(11):2551–2553.

36. Kalwarczyk, T., M. Tabaka, and R. Holyst. 2012. Biologistics--diffusion coefficients for complete proteome of Escherichia coli. Bioinformatics 28(22):2971–2978.

37. Seabold, S. a. P., Josef. 2010. Statsmodels: Econometric and statistical modeling with python. 9th Python in Science Conference.

38. Fraser, H. I., M. Kvaratskhelia, and M. F. White. 1999. The two analogous phosphoglycerate mutases of Escherichia coli. Febs Letters 455(3):344–348.

39. Gameiro, D., M. Perez-Perez, G. Perez-Rodriguez, G. Monteiro, N. F. Azevedo, and A. Lourenco. 2016. Computational resources and strategies to construct single-molecule metabolic models of microbial cells. Brief Bioinform 17(5):863–876.

40. Zhao, Y. H., M. H. Abraham, and A. M. Zissimos. 2003. Fast calculation of van der Waals volume as a sum of atomic and bond contributions and its application to drug compounds. J Org Chem 68(19):7368–7373.

41. Perry, R. H. 1973. Chemical engineers’ handbook. McGraw-Hill, New York [etc.].

42. Savageau, M. A. 1995. Michaelis-Menten mechanism reconsidered: implications of fractal kinetics. J Theor Biol 176(1):115–124.

